# The somatosensory barrel cortex controls the spindle thalamocortical oscillation by frequency locking

**DOI:** 10.1101/2025.07.09.662963

**Authors:** Mattia Tambaro, Ramón Guevara, Marta Maschietto, Alessandro Leparulo, Claudia Cecchetto, Giorgio Nicoletti, Benedetta Mariani, Samir Suweis, Stefano Vassanelli

**Author notes:** **Materials and Correspondence** Correspondence and material requests should be addressed to S.V. or S.S. contributed equally (Mattia Tambaro, Ramón Guevara).

## Abstract

The sleep spindle is a characteristic oscillation typically observed in NREM sleep and anesthesia. It is generated by a closed-loop thalamocortical circuit that is allegedly contributing to thalamocortical gating, sensory processing and memory consolidation. Yet, the circuit intricacy in terms of electrophysiological neuronal properties and connectivity has so far contributed to hinder a clear understanding of its regulation and function. In this study, we experimentally demonstrate that, when driven by the somatosensory cortex, the spindle circuit behaves as a macroscopic single-frequency self-sustained oscillator. We frequency-modulated cortical inputs to the thalamocortical spindle circuit by periodic microstimulation of the barrel cortex in the anesthetized rat. Cortical spindles exhibited synchronization by frequency locking and not resonance, displaying a characteristic Arnold tongue, a hallmark of the self-sustained oscillator. With a rate model of the barrel cortex-thalamus circuit reproducing the oscillator behavior we show that frequency-locking can govern synchronization under whisking.

**Significance statement:** Temporal coordination across neurons (neural synchronization) is believed to be a fundamental mechanism underlying brain information processing. Neural synchornization generates oscillatory signals such as waxing and waning, approximately 10 Hz oscillations occurring during sleep and anaesthesia and known as brain spindles. Spindles are generated by a closed-loop circuit between thalamus and cortex but their function remains unknown. We demonstrate that, similarly to a methronome that adjusts its frequency to the one of an external forcing oscillator, the spindle circuit is controlled by the sensory cortex according to a frequency-locking mechanism. We suggest that frequency-locking within the thalmocortical circuitry represents a flexible frequency-adjustable mechanism tuning the processing of sensory inputs by neural synchronization.

## Introduction

Synchronization between neuronal populations generates brain oscillations that are believed to be relevant for information processing^1^ and to be involved in the etiology of neurological disorders and psychoses^2^.

Synchronization is defined in non-linear dynamics on the basis of self-sustained oscillation, a periodic, active process that results from the balance between the energy pumped into the system and its dissipation^3^. A self-sustained oscillator exhibits persistent activity and, when perturbed by a periodic external force, it synchronizes by frequency locking^3^ (i.e., it becomes entrained to the perturbation frequency; see Supplemantary Methods). For example, a metronome generates a self-sustaining oscillation (it has, indeed, an internal energy source) (Fig. 1a). If perturbed by an external periodic force, however, it becomes frequency-locked to the forcing oscillator according to a characteristic Arnold tongue^3–5^: the frequency range of synchronization widens by increasing the strength of the forcing frequency (Fig. 1b-c). Importantly, this synchronization mechanism substantially differs from the classical amplitude resonance, e.g., of a swing. A swing is not self-sustained and when the external push is over, it stops. Notably, synchronization occurs only when the forcing frequency coincides with the natural frequency of the swing.

**Figure 1.**
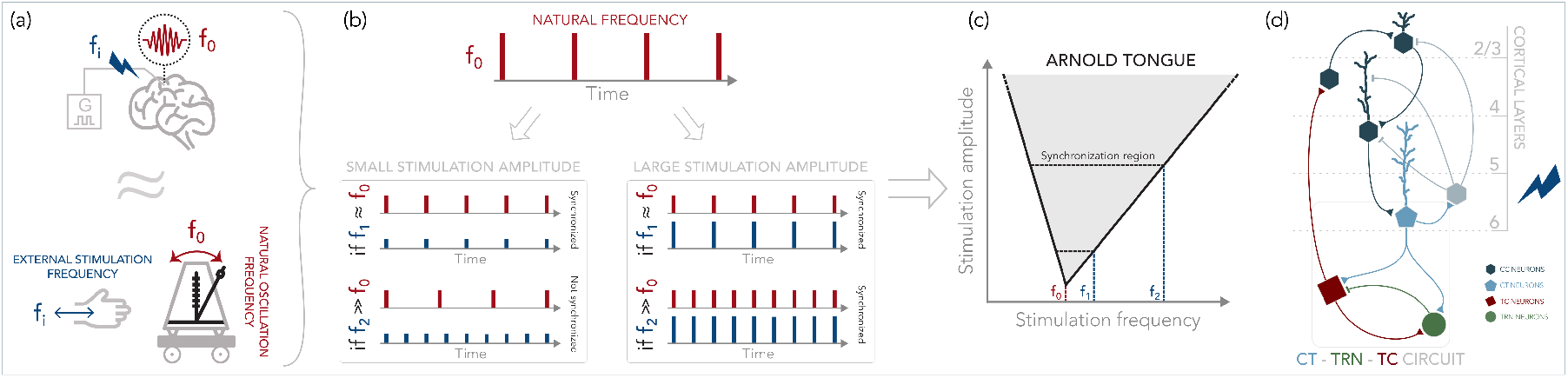
Frequency locking of a self-sustained oscillator, both in the brain and in a mechanical analogy a) Top panel: an oscillatory process with frequency *f*_0_ (in this case, a spindle in the barrel cortex of rats) is perturbed by periodic electrical stimulation at frequency *f*_*i*_. Bottom panel: A metronome represents a mechanical realization of a self-sustained oscillator. The metronome oscillates at an intrinsic frequency *f*_0_. This oscillation is self-sustained (does not vanish in spite of frictional forces) due to a source of energy. An external periodic stimulation is applied at frequency *f*_*i*_. b) When a self-sustained oscillator with natural frequency *f*_0_ (top panel) is perturbed by a small-amplitude external periodic force, the oscillator is frequency locked (synchronized) to the external perturbation if the frequency of stimulation is very close to the natural frequency of the system (bottom panel, left, upper part). Otherwise, the system is not synchronized (bottom panel, left, lower part). At large stimulation amplitudes, the system is frequency-locked to external force even if the frequency of stimulation is not close to the natural frequency (bottom panel, right upper and lower parts). c) As stimulation amplitude is increased (Y-axis), it is possible to synchronize the oscillator at a larger range of frequencies (X-axis). The region in which frequency locking can be achieved is the synchronization region or Arnold tongue (shaded area). d) Scheme of the cortico-thalamocortical loop generating the spindle oscillation with microstimulation in layer VI. CC: cortico-cortical; CT: cortico-thalamic; TC: thalamo-cortical; TRN: thalamic reticular neurons. Triangle ending: excitatory synapses. Flat ending: inhibitory synapses.

Can a self-sustained oscillator model a brain rhythm^5^? We addressed the question in the specific case of sleep spindles. Sleep-spindles are well-known waxing and waning brain oscillations of approximately 10 Hz that appear spontaneously in adults during NREM sleep and under anesthesia^6^. They are thought to be triggered by slow cortical oscillations^7,8^, while their generation and synchronization are based on recurrent thalamocorticothalamic connections. The fundamental generative circuit is a closed loop involving burst firing of inhibitory thalamic reticular neurons (TRN), leading to inhibition of thalamocortical neurons (TC), followed by post-inhibitory rebound TC firing, which re-excites RT neurons^6,9^. Cortical neurons complete the nested closed loop circuitry by sending corticothalamic projections (CT) that drive spindle synchronization (Fig. 1d). Although RT neurons are thought of as the main pacemaker, spindles are recognized as a network phenomenon involving both the thalamus and the cortex, the complexity of which is amplified by the numerous neuronal actors involved with their variegated electrophysiological repertoires^9–11^.

The function of sleep-spindles in adults is debated. Two are the major hypotheses. The first is that they contribute to govern sensory processing and disconnection within the thalamocortical circuitry to protect sleep. The second is that they promote synaptic plasticity contributing to synaptic homeostasis, memory consolidation, and learning (e.g. sensorimotor)^7,9,12,13^. Throughout development, instead, sleep-spindles have been proposed to shape the maturation of sensorimotor circuits^14,15^.

In addition to spontaneous sleep-spindles, spindle-like oscillations can be evoked by sensory stimuli (which are named alternatively evoked-spindles, evoked-afterdischarges or alpha ringing). These oscillations last for about a second after a brief sensory stimulus (e.g., a tactile stimulus) and have been found in newborn mice^15^ as well as in adult vertebrates including rats, cats and monkeys^13,16–18^. Intriguingly, evoked-spindles can be induced by sensory stimulation not only during sleep or anesthesia but also in awake or sedated animals, as demonstrated in the whisker somatosensory system of adult rats^18^ and in primates^13^. It has been suggested that sleep-spindles and evoked-spindles are generated by shared thalamocortical circuits^16,18–20^, thus playing a two-facets role for proper sensory processing in sleep and awake.

In this work, we investigated the fundamental oscillation properties of the spindle circuit when driven by cortical inputs. We used microstimulation of the somatosensory barrel cortex^18^ to frequency-modulate the cortical inputs to the thalamocortical loop of the spindle to assess its macroscopic synchronization properties. We finally used a thalamic-driven cortical firing-rate model inspired by the experimental results and predicted the modulation of the spindle oscillation by the natural whisking inputs.

## Results

We first characterized the spindles evoked in the barrel cortex by intracortical microstimulation (ICMS), and compared them with those occurring spontaneously and those evoked by single-whisker deflection. Thus, the synchronization properties of the spindle circuit under frequency-modulated ICMS (fICMS) were investigated. Electrophysiological signals (extracellular spikes and local field potentials) were acquired across the barrel cortex of rats anesthetized with urethane using a linear multi-electrode array, while performing the microstimulation in the cortical layer VI with a microelectrode of the same probe (see Methods).

### Evoked and spontaneous spindles

We confirmed that ICMS reliably evokes spindles in the somatosensory barrel cortex^18,20^ with local field potential (LFP) and spiking activity across the cortical layers following the typical waxing and waning behavior. Spontaneous, whisker-evoked and ICMS-evoked spindles showed a similar pattern (Fig. 2). The onset of spindle oscillation was at 128 ± 14.1 ms (median ± MAD) after microstimulation, preceded by a firing pause. The periodic firing epochs had a frequency of 11.6 ± 1.6 Hz (median ± MAD), were synchronous across the cortical layers, and lasted up to about one second. The duration of the firing pause after stimulation, the frequency of the oscillation, and the threshold current intensity necessary to evoke the spindle response varied slightly between trials in the same animal and between subjects. Nevertheless, spindles always exhibited the characteristic pattern. It should be noted that spindles could be evoked by single-whisker deflections only in about 25 percent of the animals (while in more than 90 percent with ICMS) and that spike synchronization across layers, initially weak, was reaching its maximum approximately 400 ms after stimulus onset (Fig. 2c).

**Figure 2.**
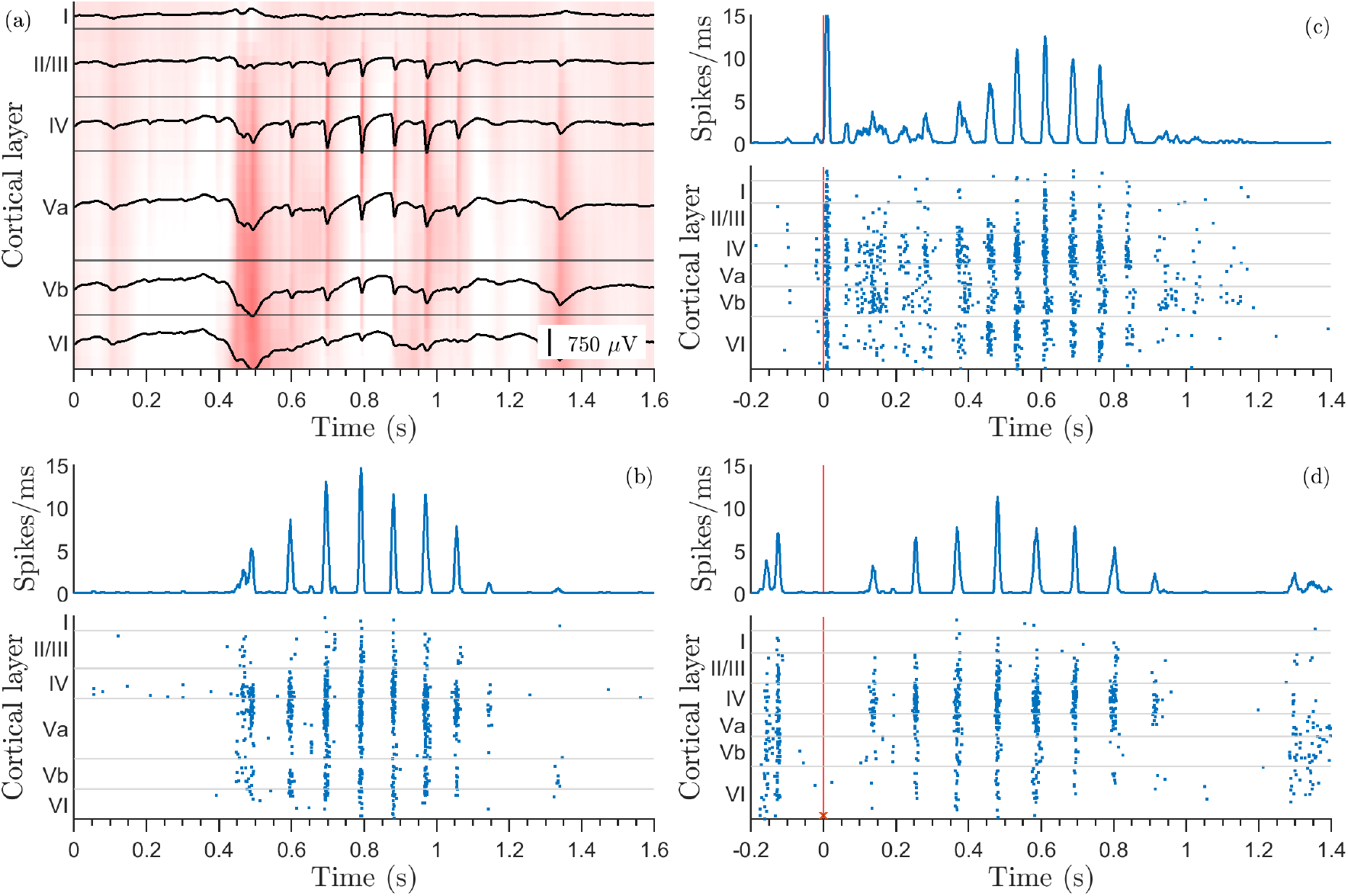
Spontaneous, whisker- and ICMS-evoked spindles. (a) LFP of a spontaneous spindle, the average LFP per layer (black trace) overlay the per-channel colormap of the activity. (b) Spiking activity of the same spontaneous spindle. Spindle evoked by a mechanical stimulation of the whisker (c), and by an ICMS pulse (d). c-d: cortical firing rate (blue trace), time and cortical depth of the spiking activity (blue dots), stimulation onset (red line) and ICMS cortical depth (red cross).

### Frequency locking of spindle oscillations

We investigated the nature of the spindle oscillation taking advantage of the fICMS approach mentioned above. Two representative spindles, recorded in the same animal and modulated by fICMS at 6 and 14 Hz are illustrated in Fig. 3a and b, respectively. Frequency-modulation across the entire 6 - 20 Hz frequency range is shown in the Supplementary information Figure S1.

**Figure 3.**
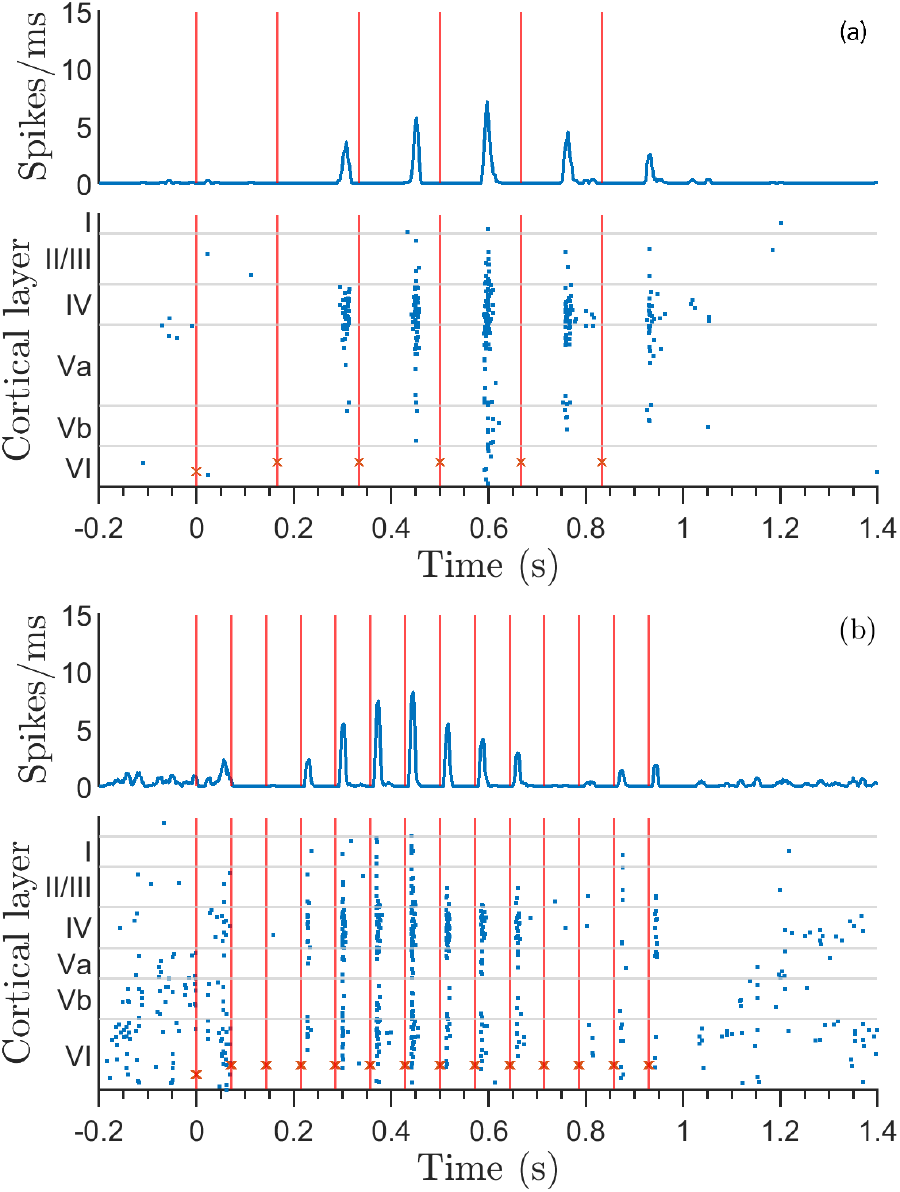
Frequency modulation of spindles by ICMS. Representative examples of spindles frequency-modulated by ICMS at (a) 6 Hz and (b) 14 Hz. Cortical firing rate (blue trace), time and cortical depth of the spiking activity (blue dots), ICMS onset (red lines) and depth (red crosses).

To assess the hypothesis that the spindle circuit behaves as a macroscopic self-sustained oscillator, we searched for the presence of frequency-locking and related Arnold tongues (Fig.4 a and b). In response to periodic intracortical stimulation, there was a slight change in the frequency of the evoked spindle oscillation. The natural frequency of the spindle, *f*_0_ (approximately 10-12 Hz), generally shifted to a different value, *f* , as a result of stimulation. If the frequency of stimulation is denoted as *f*_*s*_, then the response to stimulation can be quantified as Δ *f* = *f* − *f*_*s*_. Each trace in Fig. 4b represents Δ *f* for the entire range of stimulation frequencies at a specific stimulation amplitude. At the highest stimulation intensity (20 *µA*), Δ *f* is approximately zero over a wide frequency range, indicating that the spindle oscillation was frequency-locked to stimulation within that region. This central part of the trace is known as the “synchronization plateau”^3^, because of its flatness (Δ *f* close to zero). The synchronization plateau was approximately centered at a value corresponding to the natural frequency of the spindle oscillation ( *f*_0_ ∼ 11 Hz). Outside the plateau region, the traces had a negative slope, which demonstrates the absence of frequency locking. On the other hand, plateaus became smaller (and slightly inclined) as the stimulation intensity decreased, indicating that frequency locking was achieved within a narrower frequency range. In Fig. 4a, color-coded and interpolated Δ *f* are reported in relation to the stimulation frequency *f*_*s*_ and the amplitude (i.e., the current intensity), with the white region that indicates frequency locking. These results demonstrate that spindle oscillations are self-sustained oscillations as defined in the theory of nonlinear dynamical systems (compare to Fig. 1).

**Figure 4.**
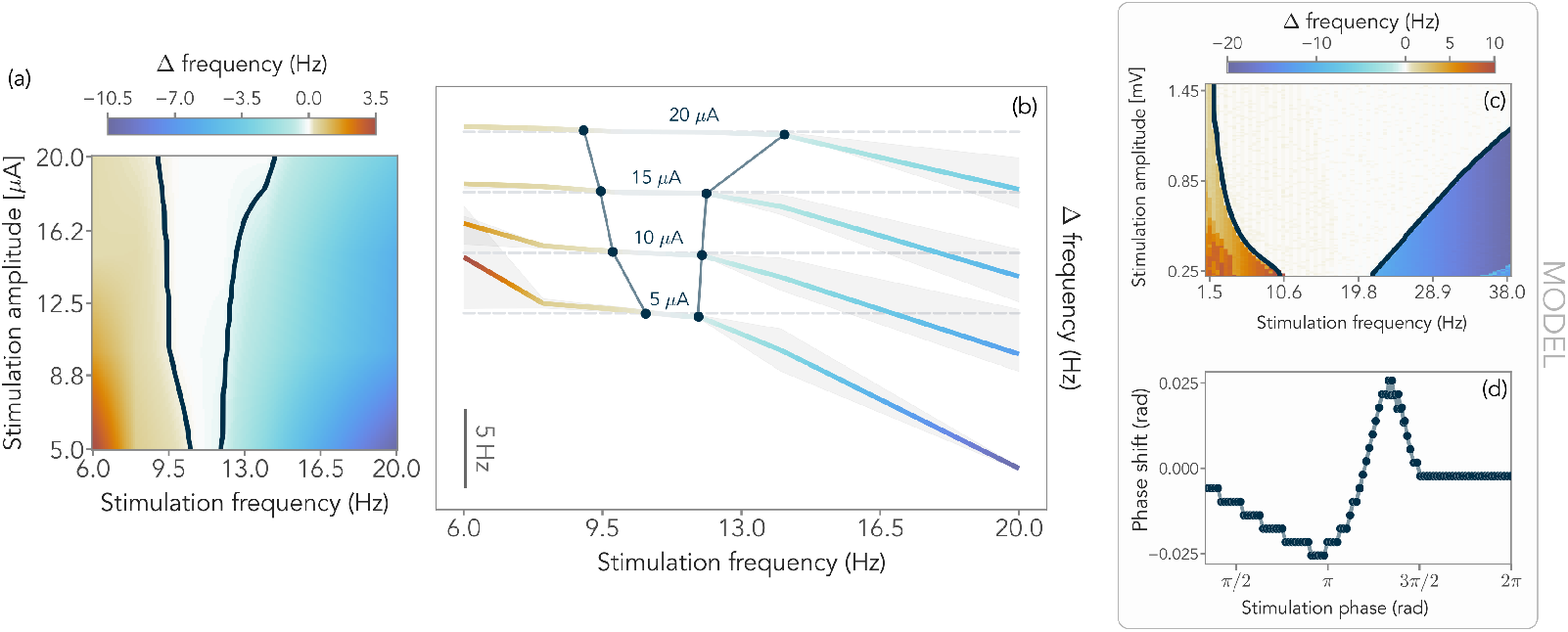
Frequency locking of spindle oscillations: Arnold tongue and synchronization plateaus. a) Arnold tongue. Trains of periodic electric pulses were applied at different stimulation frequencies, *f*_*s*_ (X-axis), and amplitudes (Y-axis) in layer VI of the barrel cortex. The frequency of the evoked spindle oscillations changed as a result of stimulation. The frequency change Δ *f* = *f* − *f*_*s*_, is color-coded and linearly interpolated. The white region is the synchronization region (Arnold tongue), at which Δ *f* = 0, (frequency locking, *f* = *f*_*s*_). In this region, the spindle was forced to oscillate at the same frequency as that of the train of pulses (black solid lines indicate the boundaries within which Δ *f* is comprised between ± 0.3Hz). b) Synchronization plateaus. Δ *f* plots as a function of stimulation frequency, *f*_*s*_, for different stimulation current intensities (median: colored thick lines; standard error of the mean: shadowed areas). In this representation, the frequency locking area is known as synchronization plateau, at which Δ *f* is approximately zero within a range of stimulation frequencies. For visualization purposes, plots corresponding to different stimulation currents are stacked and a horizontal dashed line indicates the respective Δ *f* = 0 baseline. Similarly to a), the frequency locking area (i.e., Δ *f* plateaus comprised between ± 0.3Hz), is delimited by black dots and lines. The number of stimulation trials for determining a) and b) plots are reported, for each frequency and stimulation intensity, in Table S1 of the Supplementary information. c) Arnold tongue of the mathematical model of the barrel cortex, representing frequency-locking of the spindle oscillations to periodic stimulation of the whiskers (same variables as in panel a). d) Phase response curve of the barrel cortex model.

### A firing rate model for the rat thalamus-barrel cortex predicts the emergence of spindle oscillations

We finally show that the emergence of spindle oscillations and related Arnold tongues can be predicted by a reduced model of the rat barrel cortex activity under thalamic-driven stimulation accounting for whiskers sensory inputs. This model was originally formulated by Pinto and Ermentrout (^21^) as the reduction of a spiking neural network model emulating the dynamics of the somatosensory whisker pathway^22^. This model consists of Wilson-Cowan-like equations, describing an excitatory and an inhibitory population of the barrel cortex, together with an equation describing the input from the thalamus (see the Method section for details). The parameter values of the models are kept as those obtained experimentally in the original work^22^. We only introduced a small change to the balance of feedforward (thalamic) inhibition over excitation 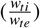 (but used values for these parameters within the standard deviations provided by^22^) to produce a periodic output, representative of a spindle oscillation in the barrel cortex.

A periodic input of the thalamus (TH) to the somatosensory cortex, representing the effect of repeated (whisker) stimulation, was modeled as a periodic sinusoidal perturbation of amplitude *A* and frequency *f* : *TH* = *A* × *sin*(2*π f t*) (where *t* is time). The stimulus amplitude and frequency were changed in the range 0.13 mV *< A <* 2 mV and 1 Hz *< f <* 40 Hz. The model has a natural frequency that is remarkably close to the frequency of the spindle oscillation, denoted as *f*_0_. In response to stimulation, the oscillation frequency is changed to *f* . A measurement of how much the system is synchronized with the external perturbation is given by Δ *f* = *f* − *f*_0_. The result of stimulation of the model at different amplitudes and frequencies is shown in Fig. 4c, representing the amount of frequency locking (Δ *f* ) as a function of the frequency ( *f* ) and amplitude (*A*) of the stimulating sensory signal. The shaded part of the graph displays the Arnold tongue. The response of the barrel cortex to mechanical non-periodic stimulation of the whiskers was also explored using the same model described above, by calculating the phase response curve (PRC) of the spindle oscillations (see Supplementary information for details). A PRC represents a complete description of the oscillator’s response to external stimulation (^3^). The PRC was calculated as follows: the model was set in the regime of parameters in which it displays a stationary oscillation. The instantaneous pulses were then delivered to the model through thalamic stimulation pulses. The pulses were delivered at random times, that is, each pulse was delivered in a different phase of the spindle oscillation. Then the phase shift induced by the stimulation was calculated as *φ* = 2*π*(*T*_0_ − *T*_1_)*/T*_0_, where *T*_0_ is the oscillation period before the stimulation and *T*_1_ is the oscillation period after the stimulation. Positive values of the shift indicate an advance of the oscillation, while negative values indicate a delay^3^. The PRC of the modeled spindle oscillation is shown in Fig. 4d. The kind of phase response curve obtained is typical of type II oscillators (PRC with advances and delays in phase, see^3^), which is consistent with the Arnold tongue obtained. Indeed, the shape of the PRC suggests that the oscillator can synchronize with frequencies that are phase-shifted (advanced or delayed) with respect to its own rhythm.

## Discussion

In this study, we have assessed how the spindle closed-loop cortico-thalamocortical circuit is responding to cortical input signals at different frequencies and amplitudes. We exploited microstimulation in the layer VI of the somatosensory barrel cortex to trigger and then to frequency-modulate the spindle oscillation. We found that the spindles frequency-lock to cortical activity with synchronization plateaus leading to an Arnold tongue. In practice, according to the typical behavior of a self-sustained oscillator, synchronization of the spindle to the cortical activity was a function of the stimulus frequency, with the synchronization frequency range widening by increasing the stimulus amplitude^3^. Within our experimental setting, the spindle oscillation was triggered with a microstimulation pulse in layer VI^18^. Right after evoking the spindle, we applied trains of pulses at different current intensities and frequencies while recording activity in cortical layers to assess neural synchronization. In conclusion, we demonstrated that, despite the complexity of the thalamic-cortical circuit underlying the spindles, its behavior is described by a well-defined, 10-12 Hz centered, self-sustained oscillator.

Following this experimental paradigm, we assumed that corticothalamic inputs to the thalamus were tuned in amplitude by changing the current intensity of microstimulation pulses. Thus, we briefly substantiate how microstimulation can tune the amplitude of corticothalamic inputs. Microstimulation in the neocortex activates primarily myelinated fibers of pyramidal neurons (mostly acting on the axon hillock or on nodes of Ranvier because of their high sodium channels density). For a current pulse of 200 *µ*s duration, the excitability constant can be as low as 300 *µ*A/*mm*^2^ for the largest myelinated fibers and as high as 27000 *µ*A/*mm*^2^ for the smallest unmyelinated axons. In practice, it can be estimated that stimulation with current intensities smaller than 25 *µ*A activated fibers (and corresponding neurons) that were roughly confined within one cortical column, and that the number of recruited fibers was approximately proportional to the current intensity^23^. It should be noted that, alternatively to the excitation of pyramidal neurons alone, a model can also be considered that assumes contextual excitation of pyramidal and inhibitory interneurons in layer VI^18^. Yet, a second different mechanism could be involved: the modulation of the firing frequency of corticothalamic neurons in layer VI, as previously demonstrated in the case of intracellular current injections^24^. In conclusion, increasing the current intensity of the microstimulation pulse strenghtens the inputs to thalamic neurons either by recruiting thalamocortical fibers or by increasing their discharge, or both.

The experimental approach we have undertaken was based on established oscillation theory. Self-sustained oscillators (even when they are noisy or weakly chaotic), respond to external periodic stimulation by changing their frequency, exhibiting frequency locking, synchronization plateaus, and Arnold tongues^3^. Finding oscillatory activity with such characteristics unambiguously defines a self-sustained oscillation. Individual neurons are known to behave as self-sustained oscillators under certain circumstances^1^. Furthermore, theoretical studies^3^ and experimental investigations (see e.g.,^4,25^) have shown that macroscopic self-sustained neural oscillations can appear as a result of the interaction of neurons in large networks. Noteworthy, this behavior fundamentally differs from that of a resonant oscillator, which responds to external forcing only when stimulated near its natural frequency, and does not exhibit frequency locking outside a narrow resonance peak. In contrast, a self-sustained oscillator can synchronize to the external stimulation also outside its natural frequency and at very low stimulation amplitudes (see Fig. 1). In other words, while resonant systems amplify inputs at specific frequencies, self-sustained systems actively adjust their dynamics to synchronize with the stimulus over extended ranges, as characterized by Arnold tongues and phase-locking regimes^3^, as shown in Fig. 4. This conceptual difference is important in terms of information transmission within recurrent thalamocortical circuits^10,11,26^ and can be extended to other neuronal and brain oscillators as well^27^. Intriguingly, we confirmed previous observations^16,18,28^ that a spindle-like rhythms can be evoked either by direct electric stimulation of the cortex, or by sensory inputs^13^ (in our case, the deflection of a single whisker) and that both responses resemble spontaneous spindles (Figure 2). This similarity is suggestive of a common circuit generator within the thalamocortical loop. What could be the role of a self-sustained spindle oscillator modulated by repeated stimulation as we observed in the barrel cortex? Evidence already exists supporting a functional role of spindle-like oscillations in awake animals, related to whisking^15,18,29–31^. In fact, there is an overlap in frequencies between whisker mechanical movements and spindles. Exploratory whisking, used by rodents to localize objects, consists of rapid movements of the whiskers in the range 5-15 Hz^32^. Spindle oscillations in the barrel cortex have a frequency range of 10 to 12 Hz. Since these two ranges overlap and since spindle-like rhythms have been observed that are coupled to the muscular swiping activity of whiskers in awake animals^15,18^, we conjecture that spindle-like oscillations could be entrained by rhythmic whisking. More specifically, oscillations could be entrained in a range that is very similar to the range of whisking, and much wider than the range of natural frequencies of spindle oscillations (again, 10-12 Hz). This is due to the nonlinear nature of frequency locking, allowing for entrainment to external perturbation even far from the natural frequency of the perturbed oscillator. It is worth emphasizing that direct electric stimulation of the barrel cortex, as realized in the experiments described here, may constitute a surrogate of the stimulation of the whiskers, as can be appreciated in panels *b* (mechanical stimulation of whiskers) and *c* (direct stimulation of the barrel cortex by a single pulse) in Fig. 2. The firing pattern across layers shows that, under anesthesia, the spindle-triggering effect of mechanical stimulation of whiskers is similar to that of direct electrical stimulation of the barrel. Both stimuli cause strong and coherent firing of corticothalamic fibers (microstimulation in particular), which is followed in the cortex by an inhibition period that is prodromal to the synchronous spindle-like oscillation. We have proposed a mathematical model that, despite its simplicity, can describe the emergent neural response of the barrel cortex to repeated thalamic stimulation. Using periodic stimulation that mimics the periodic mechanical stimulation of the whisker passing through the thalamus, we investigated the response in the barrel cortex. As observed in the experiments, we found a frequency locking of the oscillations to the periodic perturbation (Fig. 4 c), which supports the idea that the action of the whiskers on barrel oscillations is very similar to that of direct electric stimulation of the cortex. In other words, the model supports our hypothesis that spindle oscillations in the cortex are synchronized to oscillations generated by whisker activity.

Previous studies have reported phase locking between whisking and sniffing rhythms in the theta range (4–12 Hz) during rat exploration^33^, and resonant responses in vS1 neurons to whisker deflections, peaking at 4–8 Hz^34^. Similarly, spindle oscillations in the somatosensory thalamus of anesthetized rats were entrained by periodic electrical stimulation^17^. While these findings suggest a resonance mechanism in the barrel cortex, the authors did not interpret the entrainment in terms of oscillation theory, as we do here. Our results indicate a frequency-locking rather than an amplitude resonance mechanism, which requires self-sustained oscillations^3^. Evoked spindle oscillations vary in frequency across studies and conditions (e.g., 17 Hz in^18^ vs. 10–11 Hz here), suggesting a flexible mechanism for sensory information transfer from whiskers to cortex. More broadly, synchronization supports coupling and information integration in the brain^1,35^. A similar principle underlies auditory processing: hair cells in the cochlea act as self-sustained oscillators near a Hopf bifurcation, enabling high frequency selectivity and sensitivity^36–40^. Hair cell bundles can also synchronize at higher-order frequencies^41^. Analogously, our findings in the barrel cortex, along with evidence of macroscopic neural rhythms, suggest that synchronization and frequency locking may be general mechanisms in the nervous system. In summary, our study demonstrates that spindle oscillations in the barrel cortex of rats are entrained by periodic electric stimulation. We found evidence that such evoked macroscopic oscillations are self-sustained and are frequency-locked to external stimuli. The evidence is encapsulated in the characteristic Arnold tongue, typically observed when a nonlinear self-sustained oscillator is perturbed by an external periodic signal. Finally, we hypothesize that the entrainment of spindles is a key mechanism for synchronization and information transfer between the whiskers and the barrel cortex.

## Methods

### Surgical procedures

Wistar rats were housed in the Animal Facility of the Department of Biomedical Sciences (University of Padua) under standard environmental conditions. All procedures were approved by the local Animal Care Committee (OPBA) and the Italian Ministry of Health (authorization number 522/2018-PR) as described in^42^: briefly, 28 young adult rats in the age range of postnatal day 25 (P25) to P35, with the range of body weight of 80 to 160 g and of both genders were used in this study. Rats were anesthetized with an intraperitoneal induction dose of urethane (0.15/100 g of the body weight, 0.1 g/ml solution), followed after half an hour by an additional dose of the same anesthetic (0.015/100 g of the body weight)^43^ and 10 minutes later by a sub-cutaneous dose of Carprofen painkiller (Rimadyl; 0.5 mg/100 g of the body weight).

The animal was then positioned on a stereotaxic frame, and the head was fixed by teeth- and ear-bars. Body temperature was maintained at 37°C using a heating pad and monitored with a rectal probe using a homeothermic monitoring system (World Precision Instruments ATC1000 DC temperature controller) throughout the procedure. A dedicated skull window was drilled over the right somatosensory barrel cortex at stereotaxic coordinates -1 ÷ -4 AP, +4 ÷ +8 ML^44,45^. An Atlas Neuro E32+R-65-S1-L6 NT (ATLAS Neuroengineering bvba, Leuven, Belgium) probe, featuring a 32 iridium oxide microelectrodes array spaced by 65 *µ*m, was inserted into the cortex (orthogonal to the cortical surface). The Ag/AgCl reference electrode was placed near the probe and immersed in Krebs solution (in mM: NaCl: 120; KCl: 1.99; NaHCO_3_: 25.56; KH_2_PO_4_: 136.09; CaCl_2_: 2, MgSO_4_: 1.2, glucose: 11) bathing the brain.

The whiskers on the left side of the snout were first trimmed 10 mm from the mystacial pad and then individually inserted 8 mm into a 25G hypodermic needle (BD Plastipak, Madrid, Spain) glued to a multilayer piezoelectric bender with integrated strain gauges (P-871.122, Physik Instrumente, Karlsruhe, Germany) and driven by a custom-made digital closed-loop system (REF https://dx.doi.org/10.3389/fnsys.2021.709677). The whisker displacement (200 *µ*m) was triggered by applying to the piezoelectric bender a pulse of 5 ms duration through a function generator (Agilent 33250A 80 MHz, Agilent Technologies Inc., Colorado, USA). Thus, the most responsive whisker, which was providing the highest evoked LFP amplitude, was selected for recording and stimulation.

Throughout the course of the entire experiment, the level of deep anesthesia was verified by the absence of spontaneous whisking and hindlimbs and eyelid reflexes. Moreover, we monitored the presence of slow oscillations (up- and down-states) recorded by the probe during basal activity^46–48^.

### Recording and stimulation with microelectrodes

Signals were acquired at 25 kHz by a RHS 32-Channel Stimulation/Recording Headstages connected to a RHS stimulation/recording controller (Intan Technologies LLC, Los Angeles, CA, USA) and filtered in hardware by a digital bandpass filter between 0.1 Hz and 7.6 kHz. The depth of the barrel cortex layers boundaries and the electrodes assignation was based on the literature^49^ and on previous histology of the rat brain performed in our laboratory. Of the 32 Atlas probe electrodes (65 *µ*m pitch), 27 cover the entire depth of the cortex and were used for subsequent analysis. When stimulating, we run trials of 30 stimuli (6 seconds interval) for each condition (amplitude and frequency). Each stimulus consisted of a 1 second train of cathodic to anodic, symmetric bipolar current pulses (200 *µ*s duration), triggered by a waveform generator (Agilent 33250A 80 MHz, Agilent Technologies Inc., Colorado, USA). The current frequency and intensity varied from trial to trial, ranging from 6 to 20 Hz and from 5 to 25 *µ*A, as described in the Results section. The amplitude of the first pulse was set at 25 *µ*A in order to evoke spindle oscillations, while the subsequent pulses were applied from the adjacent upper channel (65 *µ*m distance). From the onset of every pulse, all the microelectrodes were grounded for 1 ms to avoid saturation of the amplifier and to minimize the stimulation artifact. Noteworthy, following this strategy, while a residual slowly decaying stimulation artifact was preventing the recording of LFP oscillations post-stimulus, spikes detection was only marginally affected. The microelectrodes pair that was selected for microstimulation was always located in cortical layer VI where we found the highest efficacy for evoking spindles (not shown). Mechanical stimulation of the whisker was performed running trials of 30 single pulse stimuli (10 seconds interval).

### Data analysis

The raw signal was filtered using a 3^*rd*^ order Butterworth filter to extract the spiking activity, with a band-pass frequency from 300 to 4000 Hz (-3 dB). The spatial average of the signals recorded in the 27 channels was subtracted from the signal at each channel to reduce correlated noise. Finally, a blanking window replaced with 0-values the filtered signal for the 15 ms following each stimulation pulse, in order to remove the sharp artifact transient and to avoid detection on the increased noise. This allowed us to exclude the false positive detection of spikes, while leaving sufficient time between train pulses (which is useful to investigate spindle activity).

The spikes were recognized by detecting negative peaks below a threshold of 4 times the median of the absolute value of the signal divided by 0.675^50^, allowing the identification of the isolated action potentials (AP) and multiunit activity (MUA). Recordings in the barrel cortex rarely showed any clear single neuron AP; the predominant activity consisted of MUA, describing a burst of spikes originating from the ensemble activity of a population of neurons with some tens of *µ*V of amplitude. This population dynamics is well identified by its firing rate, which was estimated as the sum of the spikes detected from the channels included in each layer in a moving window of 10 ms. Although this activity is noisy, its average gives information that is consistent with other measures, such as LFP.

To measure the spindle response evoked by stimulation, the cumulative spiking activity of all channels for 1 second after the stimulation onset is autocorrelated for each trial up to 250 ms shift (to find periodicity down to 4 Hz). The autocorrelation of the single trial was averaged and the result smoothed by a Gaussian window of 5 ms. The highest autocorrelation peak other than the 0-lag, if present, was compared with a threshold to find the frequency of the evoked spindle. The threshold was found by a permutation test by shuffling the position of the spikes of each trial and repeating the previous steps. Each threshold point was taken as the 95% of the maximum value reached by the average autocorrelation of the shuffled spikes for that delay. Each result was then visually inspected to check for artifacts or errors that could distort the signal and lead to incorrect detection and, in case, manually adjusted. The spindle response frequency for each stimulation frequency/amplitude pair of the subjects was found to draw the mean value and the standard error of the mean of Arnold’s tongue in Fig. 4. The response delay was instead estimated by finding the first firing rate peak after the stimuli in the spiking activity averaged per channel and trials, of experiments where spindles were recognized by the previous method. Of the 28 subjects, 3 did not show any response to stimulation at any given frequency and amplitude and therefore were discarded from further analysis.

### Mathematical model of the barrel cortex

We used a mathematical model designed to describe the barrel cortex of rats^21^. This model provides a fairly good description of the response of the barrel cortex to whisker stimulation, and is sufficiently simple so as to allow for a simulation of the barrel cortex at a mesoscopic scale. Indeed, this model is based on a biologically detailed spike-model of a single barrel previously developed by Kyriazi and Simons^22^ (with 70 excitatory and 30 inhibitory neurons, each neuron modeled as a leaky linear integrator), reduced to equations of the Wilson-Cowan type^51^. The resulting model consists of one excitatory and one inhibitory neuronal population simulating the barrel cortex. The two populations are connected with each other and receive input from the thalamus. The thalamic input is introduced in the form of time histograms (*TH*), representing the population thalamic activity. The resulting equations describe the dynamics of the average synaptic drive of both excitatory and inhibitory populations,

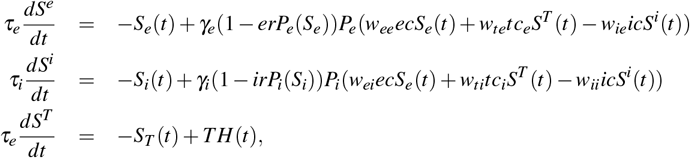

where, *S*_*e*_ and *S*_*i*_ are the synaptic drives of the excitatory and inhibitory populations, respectively. They are linked to the firing rate through the conversion factors *γ*_*e*_, *γ*_*i*_. *P*_*e*_ and *P*_*i*_ are the excitatory and inhibitory activation functions, respectively,

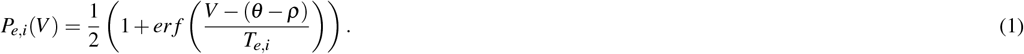

Here *V* is the membrane potential whereas *T*_*e*,*i*_ is the temperature for activation function (for both excitatory and inhibitory populations). *θ* and *ρ* are the firing threshold and the resting membrane potential, respectively. The function *er f* (*x*) is the well-known Gauss error function,

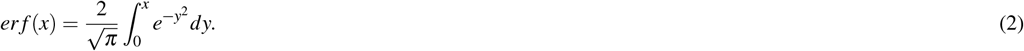

See Table 1 for a description of the parameters used in the simulations.

**Table 1.** Parameters used in the model. Their values have been inferred experimentally in^22^. Here we have changed the balance of feedforward (thalamic) inhibition over excitation 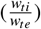 within the standard deviations provided by^22^).

| parameter | symbol | value |
| --- | --- | --- |
| resting potential (mv) | $\rho$ | -60 mV |
| firing threshold (mv) | $\theta$ | -45 mV |
| temperature | $T_e$ | 12.03 |
| - | $T_i$ | 11.62 |
| conversion factor | $\gamma_e$ | 5 |
| - | $\gamma_i$ | 15 |
| refractory parameter | $e_r$ | 2.2 |
| - | $i_r$ | 4.1 |
| connection strength (mv) | $w_{ee}$ | 1.0 |
| - | $w_{ei}$ | 1.0 |
| - | $w_{ie}$ | 2.0 |
| - | $w_{ii}$ | 1.5 |
| - | $w_{te}$ | 3.9 |
| - | $w_{ti}$ | 1.5 |
| number of synapses | $ec$ | 42 |
| - | $ic$ | 12 |
| - | $tc_e$ | 12 |
| - | $tc_i$ | 10 |
| decay constant (ms) | $\tau_e$ | 5 |
| - | $\tau_i$ | 15 |

## Data and code availability

Spiking data supporting the findings of this study and the code that analyzes it are available at doi.org/10.5281/zenodo.14537416 Raw recordings will be provided by the authors upon reasonable request..

## Acknowledgments

S.V acknowledges the financial support from the SYNCH project of the European Commission (H2020, FET Proactive, Grant ID: 824162). R.G. acknowledges the DFA UNIPD for funding through the PARD 2023 project “Response theory for brain network discovery and control”. Work by S.S. and B.M. supported by #NEXTGENERATIONEU (NGEU) and funded by the Ministry of University and Research (MUR), National Recovery and Resilience Plan (NRRP), project MNESYS (PE0000006) – A Multiscale integrated approach to the study of the nervous system in health and disease (DN. 1553 11.10.2022).

## Author contributions

R.G., M.T., and S.V. conceived and planned the experiments. M.T., M.M., and A.L. carried out the experiments. R.G., B.M., and S.S. designed the theoretical framework. M.T. analyzed the data and performed the numerical calculations. B.M. and R.G. planned and carried out the simulations. S.S. and S.V. conceived the study and were in charge of overall direction and planning. R.G., S.S., and S.V. took the lead in writing the manuscript. G.N. and M.T. devised and prepared the figures. C.C. microscopically characterized the whisker stimulation. All authors contributed to the interpretation of the results, provided critical feedback, and helped shape the research, analysis, and manuscript.

