## supplementary information for "The somatosensory barrel cortex controls the spindle thalamocortical oscillation by frequency locking"

July 9, 2025

#### 1 Supplementary Figures

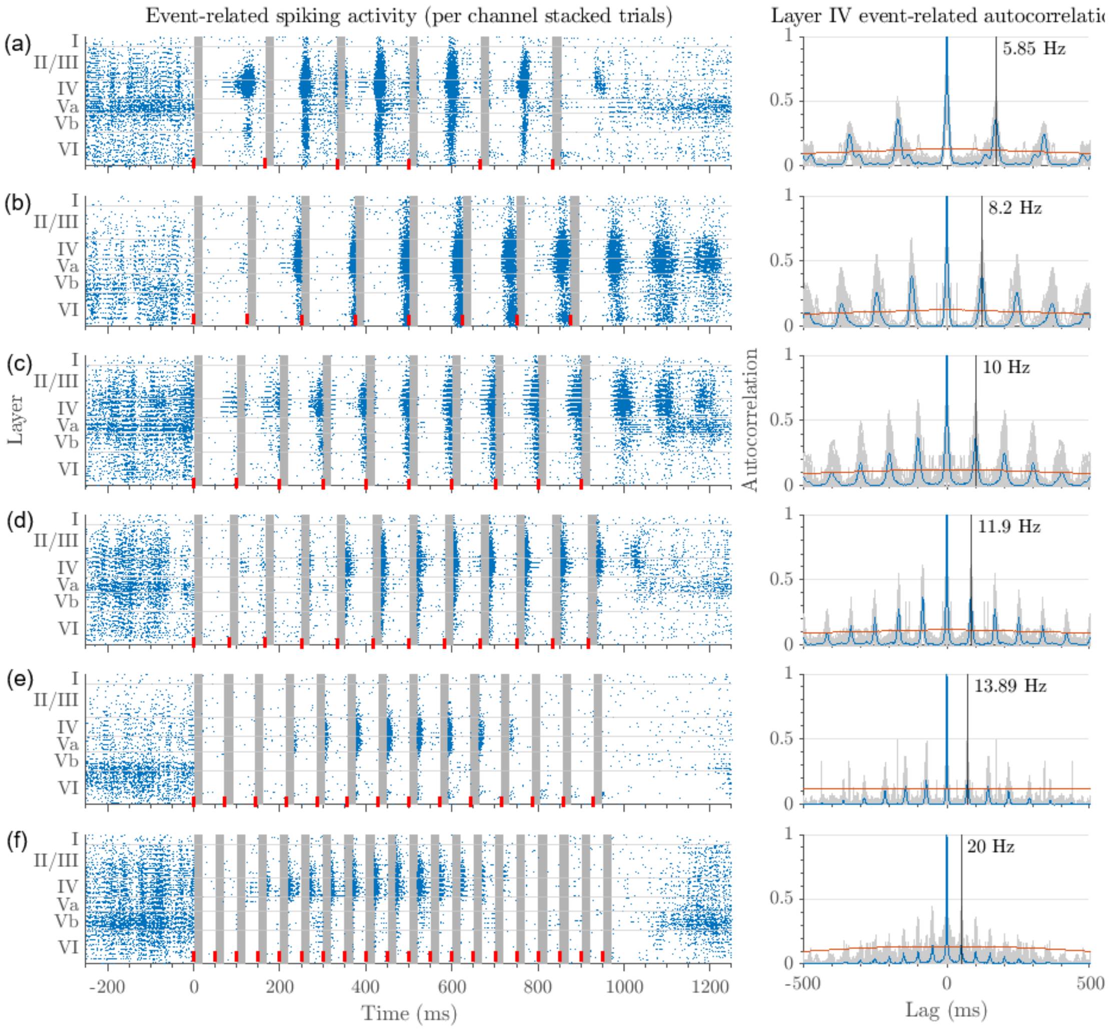


Figure S1: **Average response to different stimulation frequency.** Example rasterplots for the investigated stimulation frequencies, respectively from top to bottom: (a) 6, (b) 8, (c) 10, (d) 12, (e) 14, (f) 20 Hz. Left panels: raster plots of spiking activity in single trials (n=30) stacked for each channel (blue dots). Onset and cortical depth of the ICMSs are marked by a red bar, and the post-stimulation 'blind' window by a gray box. Right panels: spiking activity autocorrelations in layer IV (single trials in gray; average in blue). The thresholds (red lines) were computed by permutation test and the estimated spindle frequencies indicated within each panel (black line)

### 2 Supplementary Methods

#### 2.1 Synchronization

The main argument developed in the present paper rests upon concepts borrowed from the field of oscillation theory. In this brief mathematical Appendix, we clarify these ideas. In a classical context, synchronization means adjustment of rhythms of self-sustained periodic oscillators due to their weak interaction; this adjustment can be described in terms of phase locking and frequency entrainment Pikovsky et al. (2003); Izhikevich (2007).

#### 2.2 Phase of oscillations and its dynamics

A periodic oscillator whose activity is maintained for a certain period of time without damping, despite the energy loss due to friction, is called a self-sustained oscillator (to achieve this, an external source of energy is needed). In terms of dynamical systems theory, a self-sustained oscillator is a system with a limit cycle. The synchronization properties of a periodic self-sustained oscillator can be studied through the use of a special variable, called phase $\phi$, that is defined as a parametrization

| | Amplitude ($\mu$A) | | | |
|---|---|---|---|---|
| **Frequency (Hz)** | **5** | **10** | **15** | **20** |
| **6** | 60 | 420 | 540 | 390 |
| **8** | 180 | 300 | 540 | 480 |
| **10** | 90 | 240 | 390 | 210 |
| **12** | 90 | 60 | 210 | 150 |
| **14** | 90 | 60 | 210 | 60 |
| **20** | 90 | 120 | 180 | 90 |

Table S1: Number of stimulation trials for each frequency and amplitude, used to determine the Arnold tongues points

of motion on the limit cycle. The phase can always be chosen in a way that it grows uniformly in time, $\frac{d\phi}{dt} = \omega_0$ where $\omega_0$ is the natural frequency of oscillations Izhikevich (2007). The phase is neutrally stable: its perturbations neither grow nor decay. This corresponds to the invariance of solutions of autonomous dynamical systems (systems whose parameters do not depend explicitly on time) with respect to time shifts. On the contrary, the amplitude of oscillations has a definite stable value (for systems which can be described in terms of energy, this value is determined by a balance between energy influx and dissipation).

Due to the neutral stability of the phase, already a small perturbation (e.g. external periodic forcing or coupling to another system) can cause large deviations of the phase – contrary to the amplitude, which is only slightly perturbed due to the transversal stability of the cycle. Thus, with a relatively small force, the phase and the frequency of oscillations can be adjusted, without influencing the amplitude. This adjustment is the essence of the synchronization phenomenon.

#### 2.3 Synchronization by (weak) external forcing

The simplest setup for the observation of synchronization is when a periodic force is applied to an autonomous self-sustained oscillator. Stability of amplitudes and neutral stability of phases suggest the description of the effect of a small forcing in the framework of the so-called phase approximation, where only the dynamics of the phase is followed. Considering the simplest case of a limit cycle oscillator, driven by a periodic force with frequency $\omega$ and amplitude $\epsilon$, the equation for perturbed phase dynamics can be written in the form $\frac{d\phi}{dt} = \omega_0 + \epsilon Q(\phi, \omega t)$ Izhikevich (2007); Kuramoto (2003), where the coupling function Q ($2\pi$-periodic in both arguments) depends on the form of the limit cycle and of the forcing. Expanded into a Fourier series, the Q function contains fast-oscillating and slowly varying resonant terms. The latter can be gathered as $q(\phi - \omega t)$. Thus, performing an averaging over fast oscillations, the following basic equation for the dynamics of the phase difference can be obtained

$$\frac{d\Delta\phi}{dt} = -(\omega - \omega_0) + \epsilon q(\Delta\phi) \qquad (1)$$

where $\Delta\phi = \phi - \omega t$ is the difference between the phases of the oscillations and of the forcing. The function $q$ is $2\pi$-periodic. Eq.1 is called the Adler equation. On the plane of parameters of the external force ($\omega$ and $\epsilon$) there is a region $\epsilon q_{min} < \omega - \omega_0 < \epsilon q_{max}$ where Eq. 1 has a stable stationary solution that exactly corresponds to phase locking (the phase $\phi$ just follows the phase of the forcing, i.e. $\phi = \omega t$ + constant) and frequency entrainment (the observed frequency of the oscillator $\Omega = \langle \dot{\phi} \rangle$ exactly coincides with the forcing frequency $\omega$). This region is called the synchronization region, or Arnold tongue.

#### 2.4 Phase response curve

A phase-response curve (PRC) is a curve that encapsulates the synchronization properties of an oscillator by showing how its phase is altered by an external perturbation. It is obtained by stimulating an oscillator at different times ($\phi$) of its limit. The phase difference $PRC(\phi) = \phi_{new} - \phi$ (where $\phi_{new}$ is the new phase obtained after stimulation) is plotted against the original phase $\phi$. Positive (negative) values of the PRC function correspond to phase advances (delays) in the sense that they advance (delay) the timing of the next cycle. When the period of stimulation, $T_s$, is near the free-running period of the oscillator, T, the synchronization point satisfies $PRC(\phi) =$

$T - T_s$, that is, it is the intersection of the PRC and the horizontal line. Thus, synchronization indeed occurs with a phase shift $\phi$ that compensates for the input period mismatch $T - T_s$. The maxima and the minima of the PRC determine the oscillator's tolerance to mismatch. The PRC of oscillators of Class 1 Izhikevich (2007) is mostly positive, implying that such an oscillator can easily synchronize with faster inputs ($T - T_s > 0$) but cannot synchronize with slower inputs. Class 2 oscillators instead can synchronize both with faster and slower inputs.

The PRC can be used to build a phase model in the following way. Consider a dynamical system of the form:

$$\dot{x} = f(x) + \epsilon p(t)$$

describing periodic oscillations ($\dot{x} = f(x)$) , forced by a time-depended input $p(t)$. The positive parameter $\epsilon$ measures the overall strength of the input and is assumed to be sufficiently small ($\epsilon \ll 1$). What happens when we apply the same transformation to the same equation with $\epsilon \neq 0$ ? In this case the equation can be transformed into the phase model

$$\dot{\phi} = \omega_0 + PRC(\phi)p(t).$$

In other words, to synchronize an oscillator, the input pulse train must have a period $T_s$ sufficiently near the oscillators free-running period T so that the graph of the PRC and the horizontal line intersect. The amplitude of the function $|PRC(\phi, A)|$ (where $A$ is the amplitude of oscillations) decreases as the strength of the pulse amplitude $A$ decreases, because weaker pulses produce weaker phase shifts. Hence the region of existence of a synchronized state shrinks as $A$ goes to zero. This region has a tongue-shape on the $(T_s, A)$-plane, which is consistent with the Arnold tongue theory explained in the previous section Izhikevich (2007).
